# Understanding Age-Related Changes in Proprioception through Active and Passive Tasks

**DOI:** 10.1101/2023.11.06.565689

**Authors:** Erick Carranza, Sreten Franovic, Amy Boos, Elvira Pirondini

## Abstract

Voluntary control of motor actions requires precise regulation of proprioceptive and somatosensory functions. While aging has demonstrated to decline sensory processing functions, the effect of age on proprioception remains less clear. Indeed, previous studies contradict on whether passive proprioception (i.e., during externally driven movements) is affected over age. Moreover, no studies explored age-related modifications in active proprioception (i.e., during voluntary movements). Understanding these changes is critical to identify and prevent possible causes of reduced movement performances and mobility, in particular in age-related neurological conditions such as stroke or Parkinson disease. Here, we first refined a robotic protocol to assess upper-limb active proprioception and demonstrated the robustness and reliability of outcome measures over sessions. We then compared performances between young and elderly subjects during active and passive proprioceptive tasks. We found that while the two populations of subjects performed similarly in passive proprioception, elderly subjects’ acuity declined during active proprioception. Importantly, we confirmed that the decline could not be explained by impairments in voluntary motor control. Our data and robotic protocols further elucidate the debate over proprioception in elderly subjects and provide evidence that robust proprioceptive assessments should be an essential measurement in clinical settings.

**New & Noteworthy:** Upper limb position sense has been previously evaluated using passive matching protocols with the contralateral arm. These studies have found contradicting results regarding a possible decline in passive proprioception over age. Here we replicated this passive protocol and compared it to its active counterpart. Our results suggest that aging has an effect only in the active condition, leading to the hypothesis of a potential role of the cortico-spinal and cerebellar pathways in proprioception decline.

## INTRODUCTION

Voluntary movement relies on the integration of information between cognitive, motor and sensory centers within the central nervous system. Indeed, sensory feedback provides accurate internal (own movement) and external information to higher level areas in the brain that shape an adequate motor response. As we age, our body suffers from alterations that affect these functions. For instance, older adults overestimate their force production, showing evidence of sensory attenuation both in the hand and arm (1). Also, upper-limb sensory processing ability significantly decline over time in particular in passive tasks, i.e., when subjects are tasked to recognize different geometric shapes when their limb is moved passively through a path, as compared to active tasks, i.e., when subjects are instructed to use their arm actively to explore a path and to report the shape reproduced (2). Altogether, disruptions in sensory processing and integration due to aging can affect movement quality, severely impacting self-care independence, safety, mobility and balance (3–9).

Sensory integration relies on an important component for voluntary movement: proprioception. Proprioception is described as the non-visual perception of our limbs in space due to the integration of proprioceptive information provided by soft tissues like muscles, tendons and ligaments. Sensory organs known as proprioceptors sense the elongation of these tissues, providing this information to the central nervous system through sensory nerve endings and generating the sense of position (10, 11). Proprioception is a crucial component of voluntary motor control, since it provides an updated schema of the body and its loss can substantially affect motor behaviors (12, 13).

The sense of position can be investigated from a passive or active perspective. Passive proprioception refers to the identification of the limb position during an externally induced displacement, while active proprioception involves volitional movement. Only a few studies have explored the differences between active and passive proprioception, yet obtaining inconclusive results. Indeed, while previous works showed that participants discriminated more accurately at a target location during active reaching than passive movements both in healthy young subjects and in pathological conditions (14–16), Capaday and colleagues (17) did not find any difference between the active and passive conditions when subjects reached the same location. Additionally, no studies so far compared changes of these conditions with age.

Previous studies have shown that muscle spindle biology changes through life, i.e., muscle spindles decrease in number, diameter, and sensitivity (18–24). However, whether these biological changes lead to a decline in proprioception is still unclear. Indeed, using an upper-limb position matching paradigm in a cohort of participants ranging from 18 to 90 years old, Herter et al (25) found that performance in passive proprioception declined with age. Yet, recent replications of these findings showed that the effects are minimal and require a large population to be captured (1, 2, 26). Additionally, for lower-limb, a recent study reported that age has no effect on ankle passive proprioception, contradicting previous conclusions (27).

Here we aimed to tacit this debate about the differences in position sense between passive and active conditions particularly in the case of elderly subjects. For this, we built on previous passive robotic-based upper limb tasks (25, 28, 29, 29, 30) and developed an active counterpart. After validation of our protocol in young healthy participants, we recruited two populations of unimpaired participants, a young group and an older group, and we compared their performances providing novel evidence about age-related changes in the sense of position. Importantly, our experimental protocol could be used in clinical settings to assess deficits in active and passive proprioception in neurological conditions.

## METHODS

### Participants

20 young adult participants ranging from 18 to 35 years old (Mean: 28 ± 3.71 y.o., Females: 12), and 15 older participants in the range between 65 to 80 years old (Mean: 66 ± 3.86 y.o., Females: 9) were recruited for the study. The 10-item version of the Edinburgh handedness questionnaire (31) was assessed to evaluate hand dominance. Participants were considered left handed if the score was less than -40, right handed if the score was more than +40 and ambidextrous if scoring between -40 and +40. All participants scored more than +40, i.e., all participants were right handed. The exclusion criteria included history of neurological disorders or psychiatric disease or cognitive impairments as well as history of skeletal diseases that impaired elbow and shoulder joint range of motion and muscle strength. The study was approved by the University of Pittsburgh Institutional Review Board (Protocol STUDY20040260). All subjects provided written informed consent before their participation and the recordings were carried out in agreement with the Declaration of Helsinki.

### Robotic Platform

All experimental paradigms were implemented using the KINARM Upper-Limb Exoskeleton platform (**Figure 1A**, BKIN Technologies Ltd., Kingston, ON, Canada) and programmed using MATLAB-Simulink (MathWorks, Natick, MA). This bimanual robot allows movements in the 2D plane, providing gravity support. The robot works along an augmented reality screen that provides visual feedback to the user. This platform records hand-based kinematic data and elbow and shoulder joints at a frequency of 1000 Hz. For all the tasks, participants were seated straight, a seat belt restrained any compensatory movements from the shoulder and the robot was adjusted to the participant’s arm length. A cloth at the shoulder height was also used to occlude the arms during the proprioceptive tasks.

**Figure 1.**
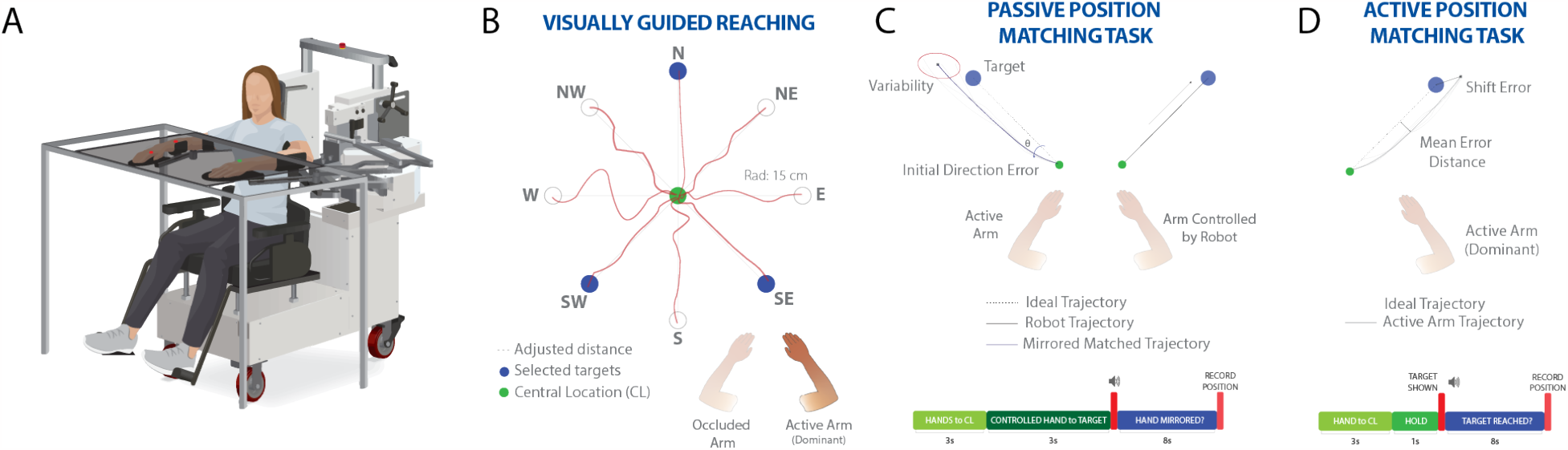
Robotic tasks. **A)** Kinarm Exoskeleton Robot. **B)** Visually guided reaching task. Participants were instructed to reach different targets located in a circular pattern. A small white dot provided visual feedback of the index location. After 3 repetitions on each target, the three targets that showed the straightest trajectories were selected for the next tasks. **C)** Passive test, i.e‥, passive position matching task. The robot controlled the non-dominant arm while the participant had to perform the motion with the dominant arm. With both arms occluded, the robot positioned both arms in the initial position. The robot moved the non-dominant arm and the participant was instructed to perform a mirror-matching movement to reach the final destination of the controlled arm. **D)** Active test, i.e., active position matching task. With both arms occluded, the participant used the dominant hand to reach a target. When the participant felt the target was reached, the final location was saved.

### Experimental Paradigm

#### Visually Guided Reaching task

The visually guided reaching task (VGR) (32, 33) was performed to obtain a baseline of the participant’s motor capabilities (**Figure 1B**). A small white dot representing the index fingertip was displayed at all times as feedback for the participant. Additionally, the participant had full vision of both arms. In each trial, the participant’s arm was positioned in a central target by the robot, locking it in place. The position of the central target was located to have the shoulder at 30°of horizontal abduction and elbow at 90°of flexion. This central target position was the same in all the tasks. An auditory cue was presented at a random delay between 100 to 700 ms, following which the robot released the participant’s arm to start the movement. The participant was instructed to reach a peripheral target located 15 cm far from the initial target as straight as possible and at a self-pace speed, i.e., as he/she would do in a regular reaching scenario. When the peripheral target was reached, the robot positioned the arm back in the central target. 8 targets arranged in a clock-like fashion were tested. Each target was presented three times during the task. After the task was completed, the three targets with the straightest trajectories were selected for the following tasks. This selection ensured that the performances reported in the following tasks (i.e., passive and active position matching) would not be influenced by the inability of the subjects to actively reach the tested targets and, thus, establishing a method applicable to individuals with motor deficits consequently to neurological disorders.

#### Passive Position Matching task

We implemented a slight variation of the passive position matching task (PPM) previously developed by Dukelow et al (28) (**Figure 1C**). Indeed, in our protocol we wanted to test the same targets for all tasks (VGR, PPM, and active position matching). Vision of both participant’s arms was occluded. For each trial, the robot positioned both arms in the central target position (i.e.,same central target of the VGR task), i.e., both arms had the shoulder at 30°of horizontal abduction and the elbow at 90°of flexion. While both arms were in the central target, the small white dot representing the index fingertip was displayed for 500 ms in order to avoid lack of visual feedback as a confound factor. Indeed, previous studies showed that, in the absence of visual feedback of the moving limb as in our case, reaching movements are more accurate and endpoint errors are smaller when vision of the starting hand position is available, compared to when such information is removed (34–43). After 500 ms, the white dot disappeared and the robot moved the participant’s dominant hand (right hand) to one of the three selected target locations (i.e., targets with the straightest trajectories in the VGR task). Once the hand was positioned in the selected target and the arm was static, a sound and visual cue was presented at the side of the non-dominant hand (left) at a delay between 100 -700 ms. The participant was instructed to move the non-dominant hand to mirror match the target position of the dominant hand. As soon as the mirror position was reached, the subject notified the experimenter and the final hand and arm positions were recorded. Next, the robot moved both arms back to the central target position to start a new trial. The position matching was repeated five times for each of the three targets.

#### Active Position Matching Task

We built on previous works (14, 16) where participants have to perform volitional upper limb motions to reach targets without visual feedback of the arms. Subjects were instructed to use the dominant hand (right) to perform the active position matching task (APM) (**Figure 1D**). As for the PPM task, vision of both participant’s arms was occluded and the robot positioned both arms in the central target position, i.e., both arms had the shoulder at 30°of horizontal abduction and the elbow at 90°of flexion. An auditory and visual cue with a random duration between 100 - 700 ms was then presented at the side of the dominant hand (right) and the small white dot representing the index fingertip disappeared. One of the three selected target locations (i.e., targets with the straightest trajectories in the VGR task) appeared on the screen and the robot unlocked the participant’s arm. The participant was instructed to perform a single as straight as possible motion to reach the target at a self-paced speed. The participant orally reported when he/she perceived that the target was reached and the experimenter recorded the final hand and arm position. Next, the robot moved the arm back to the initial position, the white dot fingertip feedback was displayed for 500 ms to remove visual bias and the participant was ready to start the next trial. Each target location was repeated five times.

### Data Analysis

The variables obtained by the VGR task provided information about motor control capabilities of the participants. These variables were: 1. *Mean distance*, which is calculated as the mean Euclidean distance between the actual trajectory and the ideal trajectory (straight line between initial and final target) and which provided a metric of movement accuracy; 2. *Trajectory smoothness*, which was calculated as the number of peaks of the mean velocity profile; 3. *Angle deviation*, which was determined as the absolute angular deviation at peak hand speed between the real and the ideal trajectory and provided a measurement of sense of direction; 4. *Maximum speed*, which was calculated by obtaining average peak speed during movement. These variables were calculated for each target and repetition separately.

For the PPM task, positional data were mirrored across the x-axis to compare location or movement of the active arm to that of the robotically-moved passive arm. We then assessed impairments in position sense using the following metrics: 1. *Shift error:* the absolute mean error distance between the end-point position of the limb moved by the robot and the subject’s opposite arm; and 2. *Variability*: the standard deviation of the averaged end-point final positions reached by the participant.

For the APM task, we estimated the active counterpart of the PPM variables. Specifically, dynamic position sense was estimated with the following metrics: 1. *Shift error*: the absolute mean error distance between the end-point position of the active arm and the target location; and 2. *Variability*: the standard deviation of the averaged end-point final positions reached by the participant. Importantly, both for PPM and APM, *Shift error* represents the accuracy of the participant in sensing the position, whereas *Variability* represents their precision in acuity.

### Statistics

All statistical comparisons of means presented in this manuscript were performed using the bootstrap method, a nonparametric method of creating resampled empirical confidence intervals for outcomes of interest, while making no assumptions of distribution of the data. For each comparison, we constructed 10,000 bootstrap samples and calculated the difference in means of the resampled data. A 95% confidence interval for the difference in means was obtained by identifying the 2.5th and 97.5th quantiles for the resulting values. The null hypothesis of no difference in the mean was rejected if 0 was not included in the 95% confidence interval. If more than one comparison was being performed at once, we used a Bonferroni correction by dividing this alpha value by the number of pairwise comparisons being performed. For our validation analysis, repeated measures t-tests were performed to analyze any impact of time on assessment outcome measures. Alpha was set to 0.05 for all analyses. All values are reported as Mean ± Standard Error. All analyses were performed using custom scripts created in Matlab 2022a (MathWorks, Natick, MA).

## RESULTS

### Performances In The Visually Guided Reaching Task Were Comparable Over Age

The visually guided reaching task allowed the assessment of motor control capabilities for both cohorts. Participants were able to reach all the target locations without difficulties (**Figure 2A**), in particular for the targets in the upper quadrant towards the left direction (Target N°: 2, 7, 8) (**Figure 2C**). The difference in trajectories across targets could be due to mechanical constraints introduced by the KINARM robot and justify our choice to perform proprioceptive assessment only on those targets that could be reached with a straight path.

**Figure 2.**
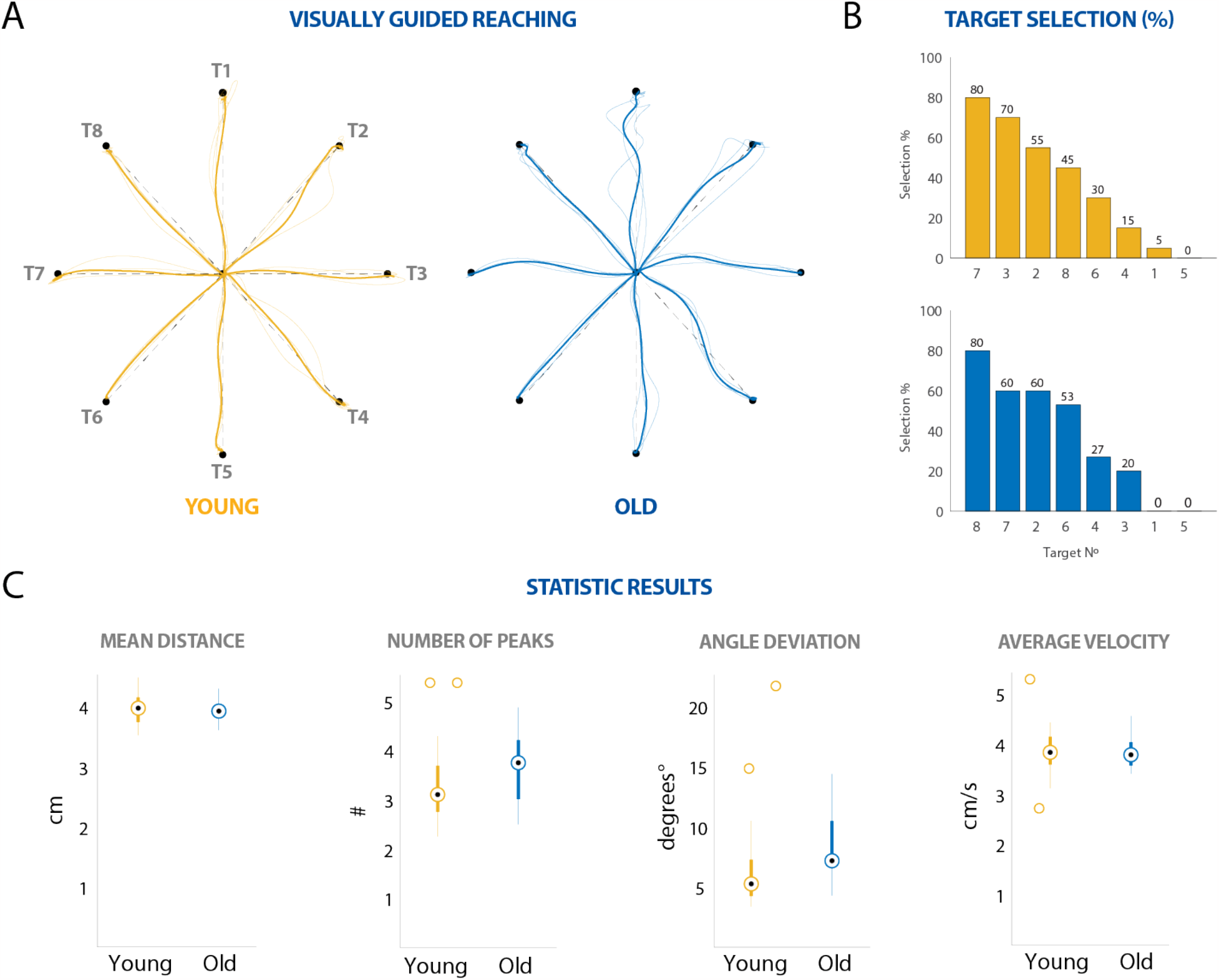
Visually guided reaching task: Young vs Old. **A)** Example trajectories for a young (*left*) and old (*right*) participant. **B)** Target selection percentage, in descending order, depicting most frequent target locations used, based on the VGR task analysis for young participants and old participants. **C)** Mean distance, number of peaks, angle deviation and average velocity were calculated to for differences between both the two cohorts. No significant differences were found between the two groups.

Importantly, older subjects did not show impairments in voluntary motor control (**Figure 2B**). Indeed, whereas movement smoothness was higher (i.e., less peaks in the speed profile) for the younger group as compared to the older group (Young: 3.1 ± 0.19; Old: 3.75 ± 0.18) and older participants showed slightly enhanced deviation as the movement started (Young: 5.26 ± 0.99°; Old: 7.17 ± 0.82°), these trends were not statistically significant. Also the mean distance was not significantly different between cohorts (Young: 3.98 ± 0.06 cm; Old: 3.93 ± 0.04 cm). Importantly, these results could not be explained by a reduced average velocity in the older subjects, which might induce more accurate movements, nor by the range of target positions (44). Indeed, the average velocity was comparable between both groups (Young: 3.83 ± 0.13 cm/s; Old: 3.78 ±0.08 cm/s). Additionally, both groups produced straighter movements when reaching towards target 7, which was located in the left horizontal plane, and target 2, which was located in the top right diagonal direction. Only the third target was different between the two cohorts. Indeed, younger participants had straighter movements towards target 3, which was located on the right horizontal plane, whereas older adults towards target 8, which was located in the top left diagonal direction (**Figure 2C**).

### Active and Passive Proprioceptive Tasks Were Reproducible Across Sessions

To demonstrate the robustness and replicability of our robotic tasks and variable measures we performed two tests. First, we assessed whether the variable measures were consistent over time. For this, five young participants performed the tasks approximately a year after the first assessment (430 +/-12 days). In a group analysis, there were no significant differences in *Shift Error* between sessions both for PPM and APM (PPM task [S1: 3.61 ± 0.86 cm | S2: 4.6 ± 0.34 cm]; APM task [S1: 1.92 ± 0.25 cm | S2: 2.24 ± 0.18 cm], **Figure 3**). Interestingly, the group showed decreased *Variability* in S2 (APM: 0.96 ± 0.24 | PPM: 1.54 ± 0.34) compared to S1 (APM: 1.01 ± 0.11 cm | PPM: 1.96 ± 0.11 cm) for both APM and PPM, but the difference was not statistically significant. Importantly, for the APM task the robustness of the *Shift Error* was preserved also at the individual level. Indeed, only one participant showed significant differences for the *Shift Error* in the APM task across sessions; whereas three participants had a significantly different *Shift Error* in the PPM task. Interestingly, the *Variability* showed a different trend. Indeed, three participants had a significantly different *Variability* in the APM across sessions; whereas none of the participants had different *Variability* in the PPM task.

**Figure 3.**
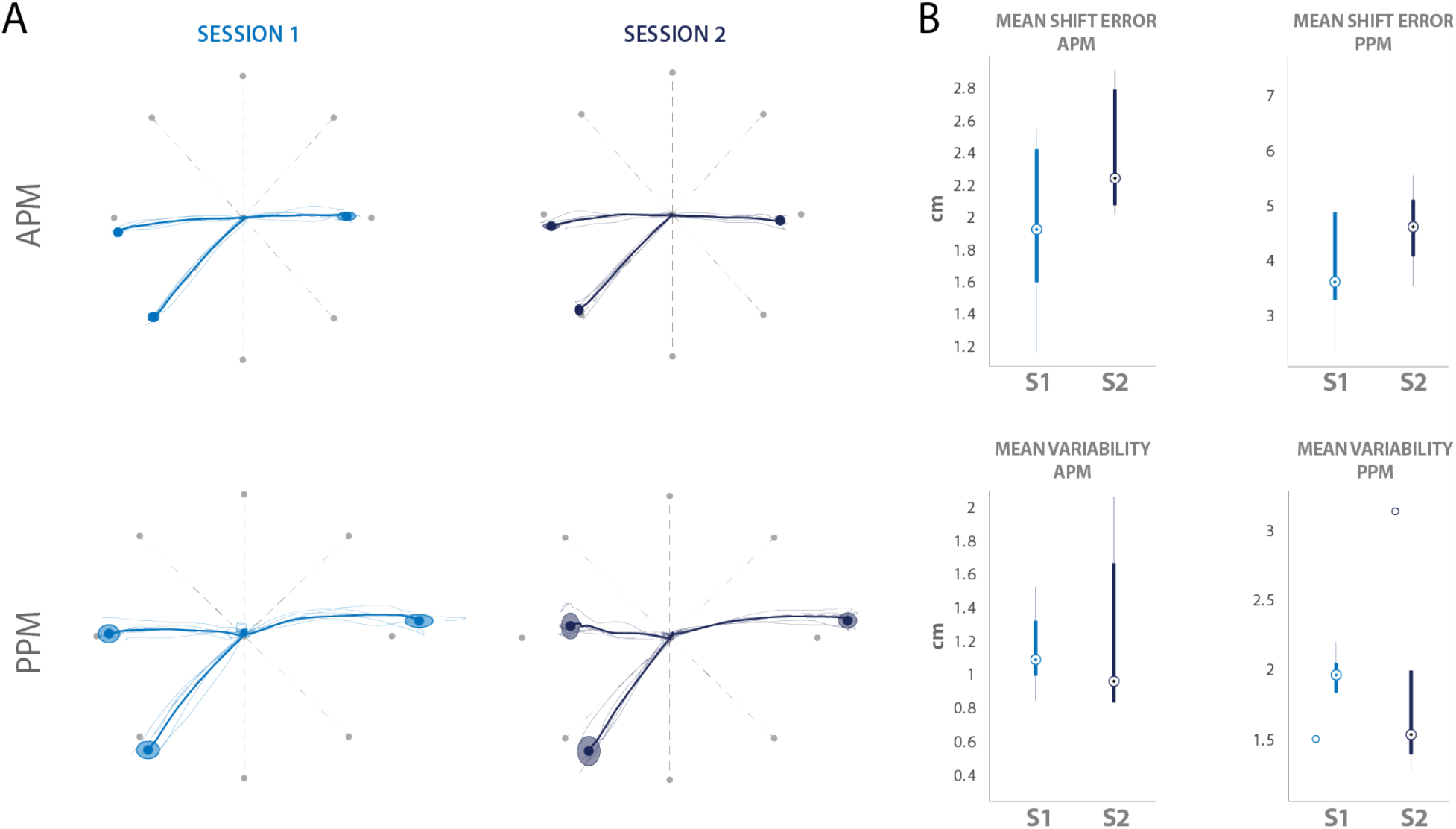
Multi-session recordings: Young participants. Five young participants performed the APM and PPM tasks a year after the first session. **A)** Example of APM and PPM performance of a subject for both sessions. The trajectories as well as the final hand positions remained consistent even after a long-term assessment. **B)** The mean *Shift Error* and *Variability* were compared between sessions both for APM and PPM. There were no significant differences between sessions.

As a second validation of the robustness of our robotic protocol, we demonstrated that the obtained results in the young and old cohorts in the PPM task were similar to those obtained in previous works (28). The KINARM platform offers the PPM task as a standard test and provides normative ranges of shift error and variability obtained over hundreds of subjects. Importantly, our results for both variables fitted inside the provided normative ranges for both cohorts (**Table 1**), demonstrating that our robotic task parallels previous tests despite the difference in target selections and the reappearance of the finger index as visual feedback.

**Table 1.** Passive Position Matching results compared to normative dataset obtained by the Kinarm platform.

| Variable | Young |  | Old |  |
| --- | --- | --- | --- | --- |
|  | Our Results<br>[range]<br>(median) | Normative<br>Range | Our Results<br>[range]<br>(median) | Normative<br>Range |
| *Shift Error* (cm) | [1.60 - 7.46]<br>(3.91) | [2.8 - 8.6] | [2.63 - 7.35]<br>(5.2) | [2.9 - 11.7] |
| *Variability* (cm) | [1.17 - 4.6]<br>2.05 | [1.9 - 5.6] | [1.24 - 10.25]<br>2.14 | [2 - 7.2] |

### Participants Perform With More Accuracy and Precision During Active Proprioceptive Tasks

After selection of the most reproducible targets, we first compared acuity performances during passive and active proprioception for the two population groups (**Figure 4**). For the young participants, we found that *Shift Error* and *Variability* were significantly higher during PPM compared to the active counterpart (PPM *Shift Error*: 4.12 ± 0.35 cm, APM *Shift Error*: 2.69 ± 0.24 cm, p<0.025; PPM *Variability*: 2.21 ± 0.16 cm, APM *Variability*: 1.60 ± 0.17 cm, p<0.025). Similar results held true for the older cohort for both parameters (PPM *Shift Error*: 5.03 ± 0.36 cm, APM *Shift Error*: 3.66 ± 0.20 cm, p<0.025; PPM *Variability*: 3.15 ± 0.64 cm, APM *Variability*: 1.590 ± 0.12 cm, p<0.025). Similarly to previous work (45–47), these results demonstrate that proprioception is more accurate and precise during active voluntary movement and that these disparities between active and passive tasks persist regardless of age.

**Figure 4.**
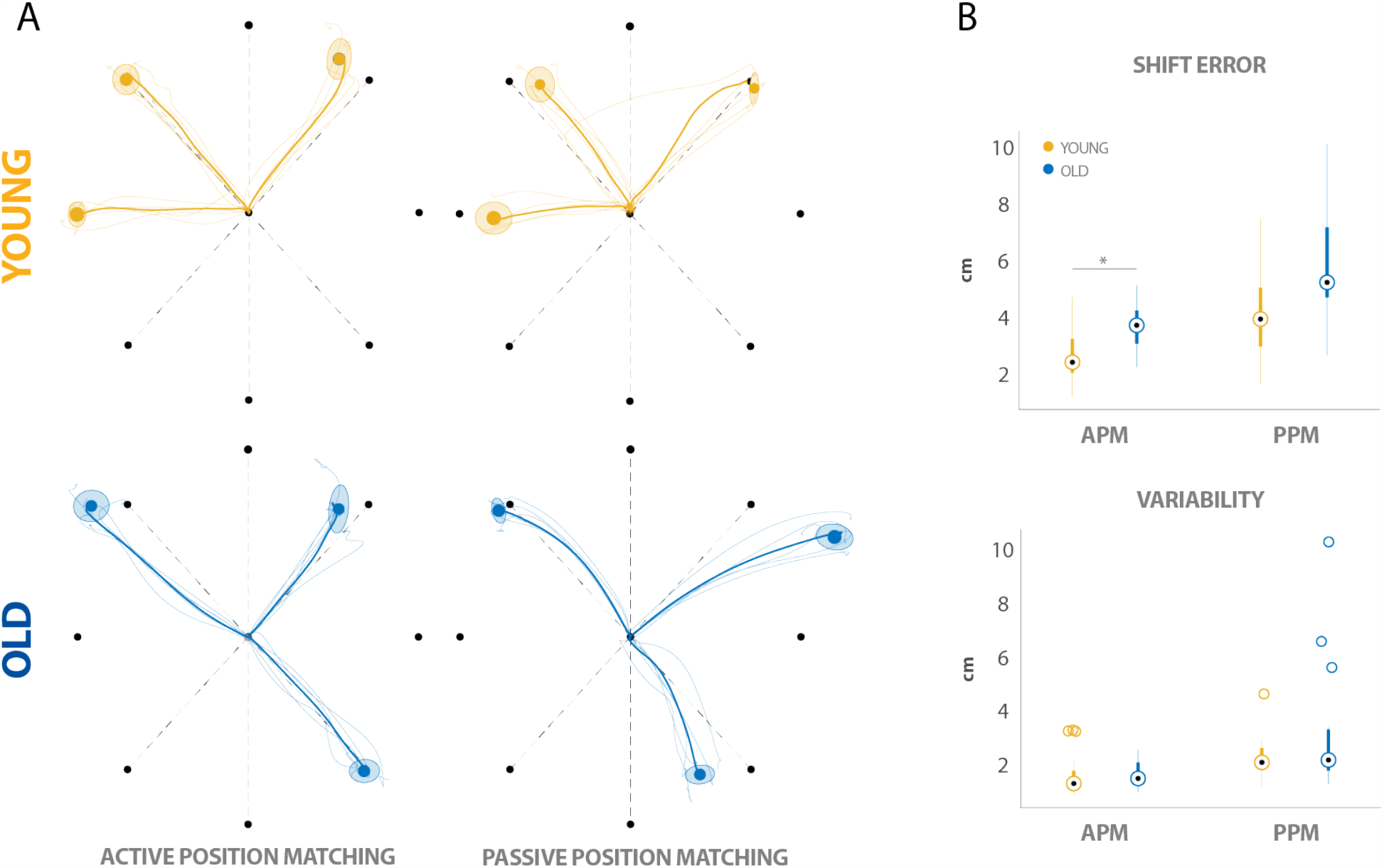
APM and PPM Tasks: Young vs Old. **A)** Examples of the participants’ performance for the APM and PPM for one young and one old participant. **B)** Boxplot comparisons of outcome measures: *Shift Error* (accuracy), *Variability* (Precision). The *Shift Error* measured during the APM task showed significant differences between the young and old cohorts. Asterisk (**\***) denotes significant differences (p<0.05).

### Aging Can Significantly Impact Accuracy of Active Proprioceptive Tasks

Next, we investigated the influence of age on the same acuity parameters, accuracy and precision, in both active and passive proprioceptive tasks (**Figure 4**). For the passive task, the older cohort performed less accurately and precisely, yet differences were not significant between the two cohorts (*Shift Error:* 5.03 ± 0.36 cm and 4.12 ± 0.35 cm for old and young participants, respectively, p>0.05; *Variability*: 3.15 ± 0.64 cm and 2.21 ± 0.16 cm for old and young participants, respectively, p>0.05).

For the active proprioceptive task, instead, results show that aging can significantly impact the accuracy of task completion (*Shift Error*: 3.66 ± 0.20 cm and 2.69 ± 0.24 cm for old and young participants, respectively p<0.025). Interestingly, active tasks did not elicit any differences among the precision of task completion between age groups (*Variability*: 1.59 ± 0.12 cm and 1.60 ± 0.17 cm for old and young participants, respectively, p>0.05).

## DISCUSSION

With the increase of life-expectancy, aging conditions are escalating so as our need to identify and treat these conditions. Disruptions in voluntary motor control due to aging can severely impact independence, safety, mobility and balance (3–9). Here we refined a robotic protocol to assess participant’s upper-limb motor capabilities and disentangle them from passive and active proprioceptive acuity. Importantly, comparison between active and passive conditions showed that precision and accuracy were higher in the active condition for both cohorts corroborating previous findings (14, 16, 45–47). This enhanced proprioception during active versus passive movement could be due to a cerebellum-dependent mechanism. Indeed, efference-copy-based predictions of body position could augment peripheral proprioceptive signals during self-generated movements (45) but not during externally induced movements (i.e., passive movements). While this difference between passive and active movements was preserved with age, elderly subjects’ accuracy declined during the active proprioceptive task suggesting that aging has a significant impact in hand position location during self-generated movements. For the passive task, instead, we did not find age as a determinant factor for decreased performance contradicting previous studies. In fact, Herter et al (25) performed the passive arm position matching task in a cohort of around 200 participants ranging from 20 to 90 years old. They found that *Shift Error* and *Variability* were affected by age. However, as also corroborated in following studies, effect sizes were small (1, 2, 26) paralleling our results.

Below, we further discuss these findings with an emphasis on the possible mechanisms underlying the changes observed with aging further elucidating the debate on proprioceptive abilities in elderly subjects. We then argue the importance of some aspects of our robotic protocol and the robustness of our outcome measures for a clinical use of these assessment protocols.

### Decline in active proprioceptive acuity: the cortico-spinal pathway

In primates, the CST is the most prominent neural pathway of voluntary motor control. CST axons originate from cortical layer V, pass through the posterior limb of the internal capsule, (a deep white matter structure in the inferomedial part of each cerebral hemisphere (48)), and terminate in the contralateral spinal cord where they form monosynaptic connections to the motoneurons. These cortico-motoneuronal connections have long been considered the most important tract for volitional control in primates, controlling skilled hand movements and manual dexterity (49-53). However, they are only one of the components of cortico-spinal innervation. Indeed, studies in monkeys have shown that CST axons innervate the intermediate zones and most importantly the dorsal horn, where sensory afferents enter the spinal cord (52–54). These innervations have been recently hypothesized to be responsible for cortico-spinal regulation of the sensory afferents (48, 55, 56). Indeed, propriospinal Chx10 interneurons, which relay information to the lateral reticular nucleus and the cerebellum, have been identified as critical for skilled reaching and receive direct input from the CST (57, 58). Additionally, the CST has direct connections with GABA interneurons in the spinal cord, which modulate sensory proprioceptive afferent terminals and produce primary afferent depolarization (PAD) (59, 60) playing a pivotal role in skilled forelimb reaching (61).

We could therefore hypothesize that age-related changes in the CST may influence its modulation role of proprioceptive and sensory information at the spinal cord level affecting active proprioceptive acuity. In this regard, age-related changes have been observed in the CST and other descending pathways originating from higher-order motor areas. For instance, Jang et al. (62) utilized diffusion tensor imaging to investigate fiber deterioration from motor cortical areas and found a significant decrease in the number of descending fibers arising from pre-motor cortex and supplementary motor area in a cohort of 70-year-old participants compared to a younger group. Additionally, Rozand et al. (63) demonstrated significantly reduced corticospinal excitability in the fibers projecting to the upper-limb muscles as a consequence of aging. Similarly, Terao et al. (64) reported a significant decrease in small myelinated projections from the CST to cervical levels of the spinal cord in aging populations. While these changes do not seem to affect voluntary reaching movements, as movement speed, accuracy, and smoothness during the VGR task was comparable between cohorts, they might influence regulation of proprioceptive information and consequently proprioceptive acuity. This effect might be stronger during self-generated movements (i.e., active proprioception) as compared to passive movements because of the stronger activation of the CST during volitional movements.

### Decline in active proprioceptive acuity: the cerebellum pathway

Another explanation of the results obtained in our experiments can be found in studies conducted with patients with cerebellar lesions. The ability to perform precise reaching movements relies on accurately estimating the current limb position and generating calibrated motor commands that are updated based on movement requirements. The cerebellum plays an essential role in movement by making precise adjustments of motor actions and is believed to be involved in the formation of internal models, both forward and inverse, which enable the prediction and correction of movement trajectories (65). While the involvement of the cerebellum in movement was traditionally thought to be solely for motor control purposes, more recent findings have revealed cerebellar projections to sensory areas in the cortex (66), suggesting a potential role in proprioception. Previous studies have demonstrated that patients with cerebellar damage struggle with active proprioceptive discrimination tasks compared to control subjects, while performing comparably well in passive versions of the same tasks (14, 67). These findings align with our observations in elderly suggesting that a cerebellar degeneration with age may contribute to the observed differences. In this regard, Bernard and Seidler (65) proposed a framework to explain how the cerebellum can influence performance as a function of age, particularly through the degradation of internal models. Aging may have a diminishing impact on cortico-cerebellar networks, as well as on cerebellar morphology and function, thus affecting the development of new internal models and the preservation of existing ones. Similarly to the CST, the degeneration of the cerebellum might not be sufficient to affect gross reaching movements as those tested on the VGR task, but it may be enough to affect precision and accuracy in active proprioception. Finally, we cannot rule out the possibilities that deficits in active proprioception might not be the results of co-occurring cortico-spinal and cerebellar changes.

### Towards a clinical use of active proprioceptive tasks

Further behavioral and electrophysiological studies in a larger population are now necessary to corroborate our findings and to demonstrate the anatomical pathways involved in the age-related changes observed in our study. However, the robustness of the outcome measure over time suggests that our robotic tasks could be used in clinical settings for the assessments of passive and active acuity. Importantly, our active proprioception test is derived by previously widely used passive proprioception tests (25, 28–30), and the same movements and target positions are assessed both for active and passive tests. This allowed us to not only compute similar measures of position sense both for active and passive tests, but also to directly compare them during the same movements in order to gather a unique characterization of sensorimotor integration. The three targets with the straightest and smoothest movement trajectories are chosen for passive and active tests ensuring that the proprioceptive performances are not be influenced by the inability of the subjects to actively reach the tested targets. This is extremely important in clinical settings with neurological patients experiencing deficits in reaching movements. Therefore, our approach will be applicable to individuals with different neurological conditions and level of motor impairments expanding our ability to assess motor control in patients.

## CONCLUSIONS

Our findings suggest that participants are more accurate and precise when performing active proprioceptive tasks compared to passive tasks. Furthermore, we found that accuracy (*Shift error*) was significantly impacted by age, while precision (*Variability*) did not show this same susceptibility to influence. Finally, both modalities of proprioceptive evaluation were deemed reproducible across sessions and thus validated for future investigations. Our findings may serve further research in a two-fold manner: as a normative baseline for comparison of impaired individuals and as a guideline for future research using age-adjusted methodology.

## Author’s Contribution

E.P. and E.C. conceived the study and designed the experiment. E.P. secured the funding. E.C. implemented the KINARM sensory tasks. A.B. and E.P. implemented patient recruitment, eligibility and monitoring and coordinated management of the study. E.C. performed the experiments. E.C., E.P. and S.F. designed the statistical data analyses. E.C. and S.F. analyzed the data. E.C. and S.F. created the figures. E.C and E.P. wrote the paper and all authors contributed to its editing.

## Acknowledgments

The study was executed through the support of internal funding from the Department of Physical Medicine and Rehabilitation at the University of Pittsburgh to EP. Additional support was provided by the Association of Academic Physiatrists Rehabilitation Research Experience for Medical Students (RREMS) and the Michigan State University College of Human Medicine MSUFCU Dean’s Choice Grant Award to SF.

